# Machine learning-based investigation of regulated cell death for predicting prognosis and immunotherapy response in glioma patients

**DOI:** 10.1101/2023.08.28.555146

**Authors:** Wei Zhang, Hongyi Liu, Ruiyue Dang, Luohuan Dai, Hongwei Liu, Abraham Ayodeji Adegboro, Yihao Zhang, Nian jiang, Xuejun Li

**Affiliations:** Department of Neurosurgery, Xiangya Hospital, Central South University, Changsha, China; Hunan International Scientific and Technological Cooperation Base of Brain Tumor Research, Xiangya Hospital, Central South University, Changsha, China; Department of Oncology, Xiangya Hospital, Central South University, Changsha, China

**Keywords:** Regulated cell death, Glioma, Machine learning, Prognosis, Immunotherapy, Immune infiltration

## Abstract

**Background:** Glioblastoma is a highly aggressive and malignant type of brain cancer that originates from glial cells in the brain, with a median survival time of 15 months and a 5-year survival rate of less than 5%. Regulated cell death (RCD) is the autonomous and orderly cell death under genetic control, controlled by precise signaling pathways and molecularly defined effector mechanisms, modulated by pharmacological or genetic interventions, and plays a key role in maintaining homeostasis of the internal environment. The comprehensive and systemic landscape of the RCD in glioma is not fully investigated and explored.

**Method:** After collecting 18 RCD-related signatures from the opening literatures, we comprehensively explored the RCD landscape, integrating the multi-omics data, including large-scale bulk data, single-cell level data, glioma cell lines, and proteome level data. We also provided a machine learning framework for screening the potentially therapeutic candidates.

**Result:** Here, we explored RCD-related phenotypes, investigated the profile of the RCD, and developed a RCD gene pair scoring system, named RCD.GP signature. Using the machine learning framework consisting of Lasso, RSF, XgBoost, Enet, CoxBoost and Boruta, we identified seven RCD genes as potential therapeutic targets in glioma and verified the SLC43A3 by q-PCR in glioma grades and glioma cell lines.

**Conclusion:** Our study provided comprehensive insights into the RCD roles in glioma, developed a robust RCD gene pair signature for predicting the prognosis of glioma patients, constructed a machine learning framework for screening the core candidates and identified the SLC43A3 as an oncogenic role and a prediction biomarker in glioblastoma.

## Background

Gliomas are the most common primary malignant tumors of the central nervous system, with approximately 10,000 new cases diagnosed each year in the United States[1]. According to the 2016 World Health Organization (WHO) classification of CNS tumors, gliomas are categorized into four grades (I-IV). Grades I and II are considered as low-grade gliomas (LGG), while grades III and IV fall under high-grade gliomas (HGG). Among these, grade IV gliomas, also known as glioblastoma multiforme (GBM), exhibit the highest degree of growth aggressiveness[2]. Unfortunately, despite advances in diagnosis and drug therapy, glioblastoma remains incurable, with a median survival time of 15 months and a 5-year survival rate of less than 5%, leading to an unfavorable prognosis. Mutations in isocitrate dehydrogenase (IDH) and O6-methylguanine-methyltransferase (MGMT) promoter methylation have a significant prognostic impact.

The only identified causative factor for GBM is ionizing radiation [2], and GBM accounts for only a small fraction of brain tumors induced by radiation. Other exposures such as cell phone use[3], cytomegalovirus[4] and germline susceptibility[5] have not been established as causative factors. The current standard treatment for GBM involves maximal surgical resection followed by radiotherapy and/or chemotherapy with temozolomide or carmustine tablet[6]. However, despite these standard therapies, glioma recurrence remains common, and patient prognosis remains poor.

The diagnosis and treatment of gliomas pose significant challenges, and RCD presents a promising area of treatment. The exploration of RCD research started with Karl Vogt’s observation of dead cells in toads in 1842, followed by the coining of the term “apoptosis” by John Kerr et al. in 1972[7]. The subsequent discovery of CED9 in mealybug development [8], and BCL2 in mammalian cells[9] triggered rapid advancements in RCD research, leading to the exploration of molecular mechanisms regulating apoptosis. In 2018, the Nomenclature Committee on Cell Death (NCCD) established guidelines for defining and interpreting cell death from morphological, biochemical, and functional perspectives. Scientists categorized cell death types into regulated cell death (RCD) and accidental cell death (ACD)[10]. ACD is an uncontrolled cell death process triggered by unexpected injury stimuli that exceed the cell’s regulatory capacity, leading to cell death. In contrast, RCD involves autonomous and orderly cell death under genetic control, governed by precise signaling pathways and molecularly defined effector mechanisms, modulated by pharmacological or genetic interventions, and plays a key role in maintaining internal environment homeostasis[10]. Major known types of RCD include autophagy-dependent cell death, apoptosis, necroptosis, iron apoptosis, parthanatos, entosis, NETosis, lysosome-dependent cell death (LCD), alkaliptosis, and oxeiptosis. The aberrant regulation of RCD has been closely linked to cancer [11], and by promoting apoptosis in tumor cells, we can potentially inhibit tumor growth and spread. Exploring the role of RCD in cancer therapy helps us understand the pathogenesis of cancer, identify key targets for controlling cell death, and develop appropriate therapeutic strategies[12].

The instability of the cancer genome allows it to accumulate a large number of point mutations during tumor development, resulting in structural alterations. The immune system plays a crucial immune-surveillance function in tumor suppression by directly killing tumor cells or triggering an adaptive immune response[13]. However, tumor cells evade immune surveillance through various mechanisms, such as defective antigen-presentation mechanisms, upregulation of negative regulatory pathways, and recruitment of immunosuppressive cell populations, which impede the effector function of immune cells and diminish the anti-tumor immune response[14]. The emergence of immunotherapy offers new hope for cancer treatment by retraining the host’s immune system and stimulating anti-tumor immune responses, including immune checkpoint inhibitors (ICIs), chimeric antigen receptor T cells (CAR-T cells), dendritic cell vaccines, and cytokine therapies[15]. These therapies improve the anti-tumor immune response with fewer off-target effects than chemotherapy and other drugs that directly kill cancer cells [16]. Immunotherapy is considered a promising strategy for treating or even curing certain cancer types; however, current clinical trials have shown that only 8% of glioma patients benefit from immune checkpoint blockade therapy[17]. Immune escape mechanisms and drug resistance in gliomas often limit the effectiveness of immunotherapy. Therefore, elucidating the mechanisms of resistance to immunotherapy could offer new possibilities to overcome the problem of low response to immunotherapy in glioma.

In this study, we conduct a comprehensive investigation of the RCD landscape of glioblastoma, analyzing both bulk and single-cell aspect. We develop a robust, accurate and RCD-related gene pair signature which has the potential to be transferable across different datasets and experimental conditions. We additionally provided a framework for screening out relatively core genes and identified SCL43A3 as a potential therapeutic oncology target.

## Methods

### Statistical analysis

All analysis was performed in R (version 4.1.3, http://www.rproject.org/). For the continuous variables and the categorical variables, the Wilcoxon rank sum test and the chi-squared test were adopted, respectively. Correlation analysis between the different variables was performed using Spearman’s coefficients. The p-value of the two-tailed tests was considered statistically significant in this study. The graphic abstract was provided in Fig 1.

**Fig 1.**
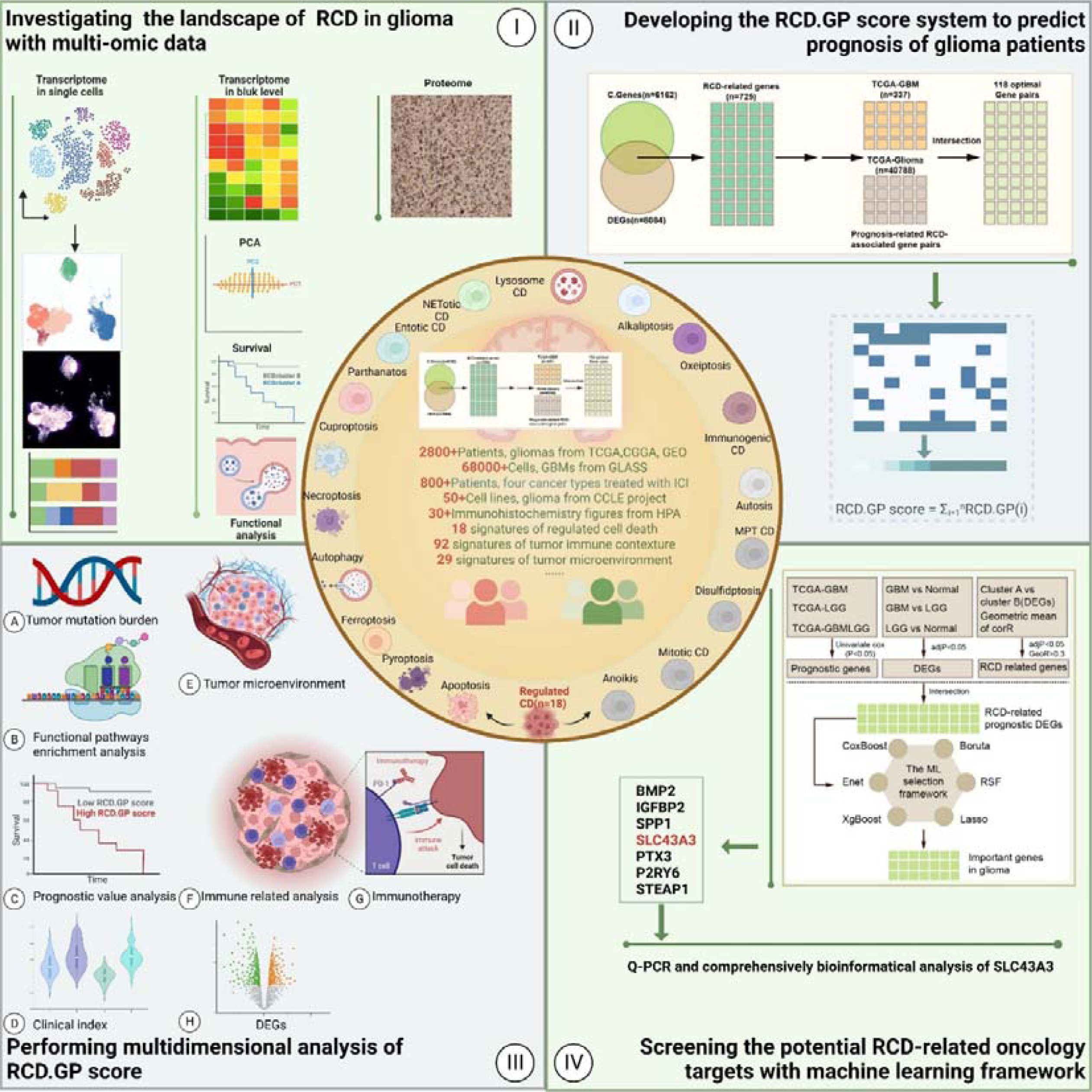
The graphic abstract of this study.

## Results

### Dysfunction of regulated cell death in glioma

Dysfunction of regulated cell death in tumors, specifically the failure of programmed cell death mechanisms, plays a significant role in tumor development and progression[11]. In normal physiological conditions, cells undergo programmed cell death, including apoptosis, autophagy, and necroptosis, to maintain tissue homeostasis and eliminate damaged or abnormal cells. Here, we comprehensively investigated the RCD level between glioma and normal brain cortex with collected 18 RCD signatures. It might seem counterintuitive that the level of RCD was dysregulated and higher in gliomas compared to normal tissue (Fig 2A). With the Spearman’s correlation analysis, we also estimated the inner regulated network among these RCD in glioma and the normal tissue (Fig 2B). The alteration of the correlation also provided solid evidence of the dysregulated RCD in glioma. This abnormal increase and dysregulation of RCD are often detrimental to the survival of patients with many types of tumors, as reported in various publications[18-20].

**Fig 2.**
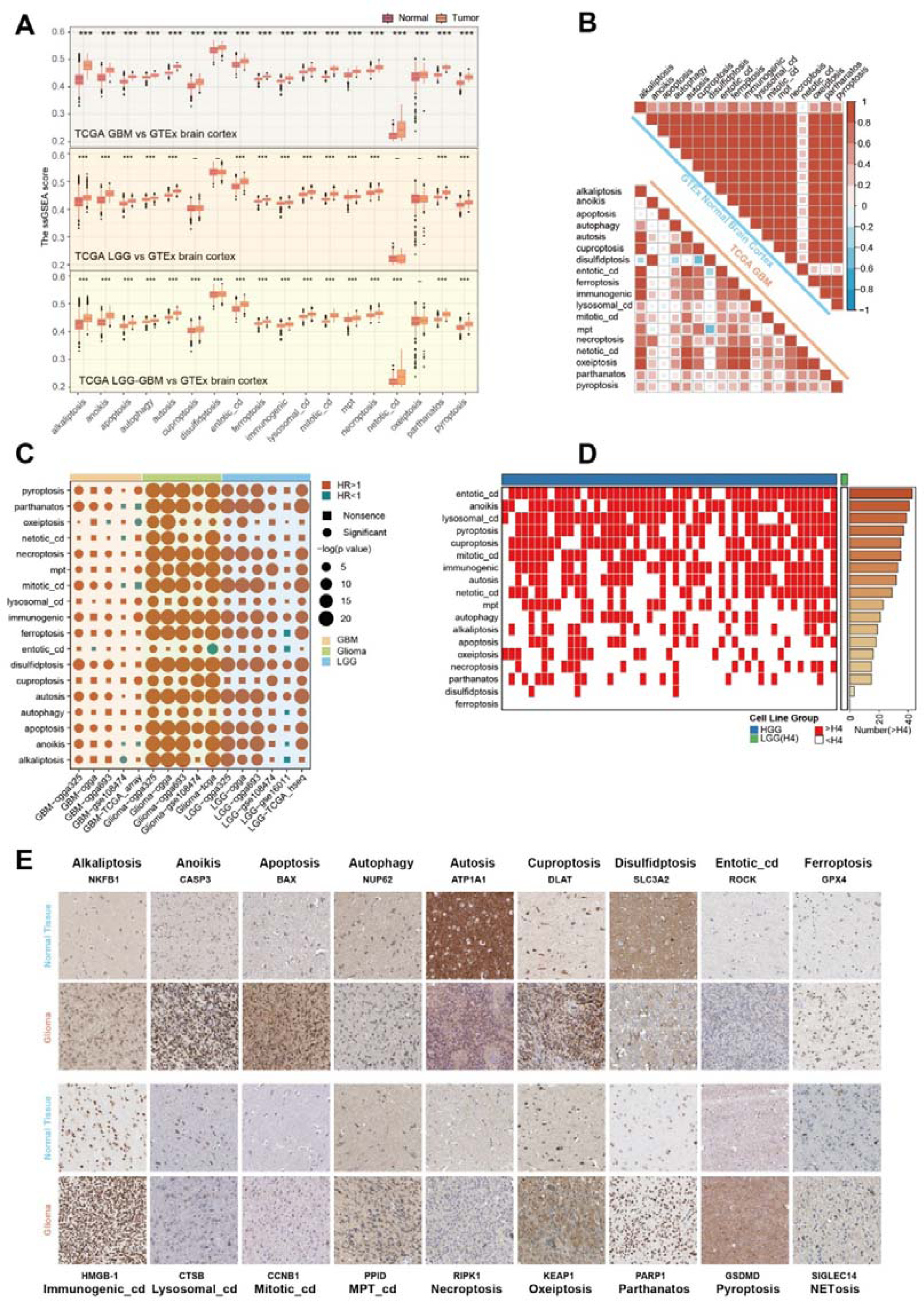
Dysfunction of regulated cell death in glioma. A. The comparison of the ssGSEA score of the 18 RCD signatures between gliomas and normal brain cortex, with the large-scale bulk transcriptomic data from the TCGA and GTEx project. The Wilcoxon rank sum test was performed. The two sided p value < 0.001 was represented by “***”. B. The spearman’s correlation of the 18 RCD ssGSEA scores in TCGA-GBM dataset and GTEx normal brain cortex dataset. C. The univariate Cox regression result of the 18 RCD signatures in datasets with LGG, datasets with GBM and datasets with glioma. The p value < 0.05 was considered as significance. D. The ssGSEA score of the RCD signatures in glioma cell lines. The red represented that the score in this GBM cell line was higher than in LGG cell line (H4), while the white represented the opposite. The right panel showing the number of the GBM cell lines in which the score was higher than in the H4. E. The immunohistochemically stained tissue sections images from the HPA of the core RCD genes indicated that the level of the RCD was different between in glioma and in normal brain tissues. The detailed clinical information of the images was provided in supplementary table 3.

To confirm whether this also applied to gliomas, we performed univariate Cox regression in several cohorts of LGG, GBM and glioma. The results revealed that the high level of the RCD was a risky factor for glioma patients (Fig 2C). The transcriptomic profiles from the CCLE project indicated that the level of most RCDs was higher in most high grade glioma (HGG) cells than in LGG cells (H4) (Fig 2D). From previous publications, we selected some core genes in individual RCDs, and immunohistochemically stained tissue sections from the HPA of these genes showed that the level of RCD was higher in glioma than in normal brain tissue (Fig 2E)[10, 12, 21-23].

Collectively, at both the mRNA and protein levels, we found a significant increase in these 18 already reported regulated modes of cell death and functional abnormalities. It is important to note that while the level of regulated cell death might be higher in gliomas, the effectiveness of these cell death mechanisms could be impaired or evaded by tumor cells, contributing to the survival and progression of gliomas.

### RCD-based patterns show distinct micro-environments

According to the consensus clustering analysis based on the profile of the RCD, the patients in the TCGA glioma dataset could be divided into two distinct RCD patterns, named RCD cluster A (n = 220) and RCD cluster B (n = 469) (Fig 3A, supplementary table 9). The PCA results indicated different characteristics of the RCD profiles between the two RCD clusters (Fig 3B). Next, we described the differences and connections between these two RCD clusters at multiple levels, including clinical features, immune microenvironment, signaling pathways and so on.

**Fig 3.**
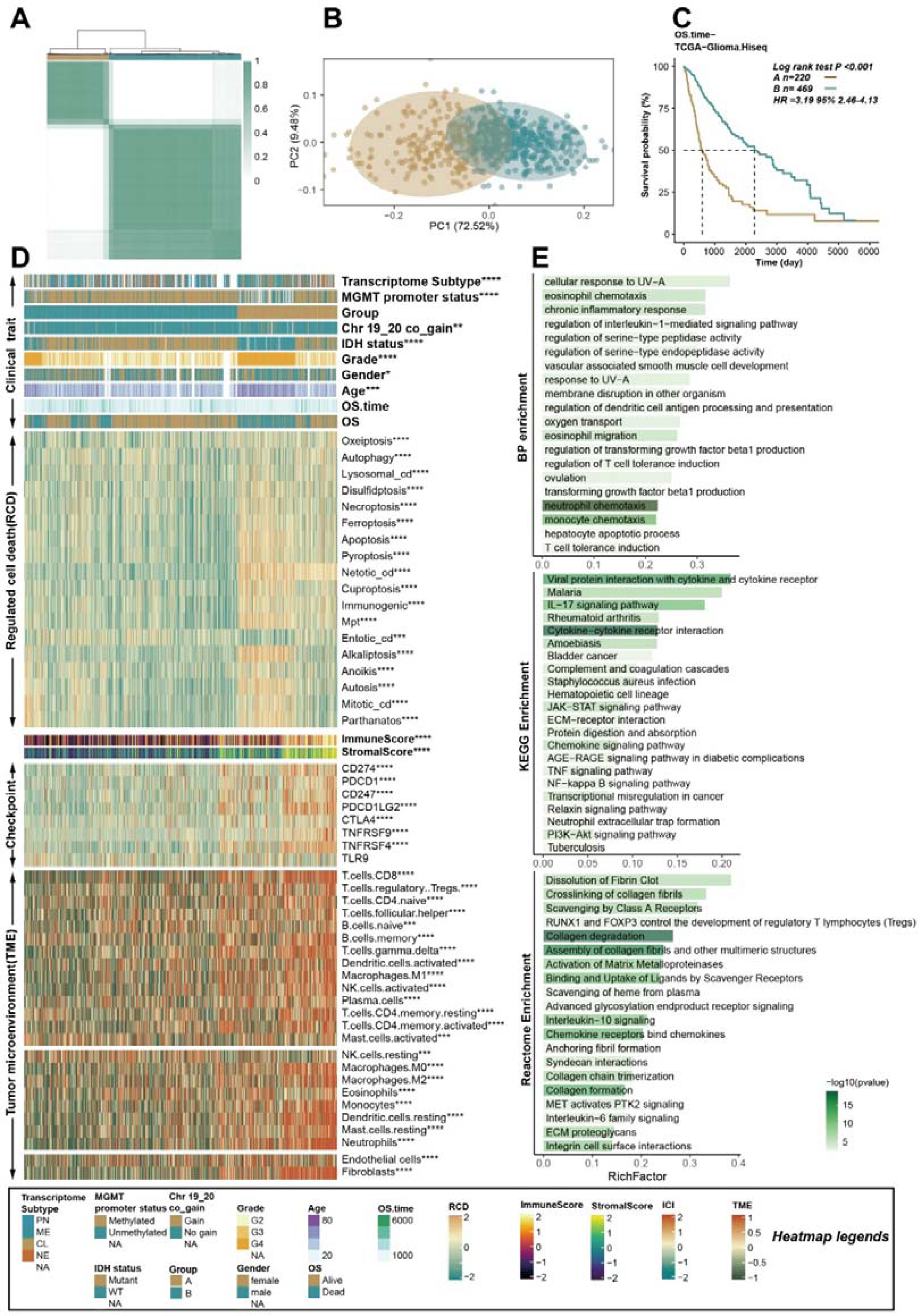
RCD-based patterns show distinct micro-environments. A. The glioma patients in TCGA were classified into two RCD clusters, named RCD cluster A and RCD cluster B, based on the consensus clustering analysis. B. The PCA analysis showing the different distribution of the RCD profile. C. The K-M curves showing that the patients in RCD cluster A had worse OS than patients in RCD cluster B. D. A heat map showing the clinical index, RCD profile, expression of immune checkpoint genes, immune score, stromal score and tumor microenvironment in the TCGA glioma cohort. The Wilcoxon rank sum test or the chi-square test was performed the assess the difference between the RCD cluster A and RCD cluster B. “*”, “**”, “***”, and “****” represented that the p value < 0.05, 0.01, 0.001, and 0.0001. E. The enrichment analysis of the DEGs between the two RCD clusters. KEGG, GO BP and REACTOME databases were included for the functional pathway analysis.

The patients in RCD cluster A had significantly better overall survival than patients in RCD cluster B (Fig 3C, log-rank test p < 0.001, HR:3.195, 95%CL:2.46-4.13). Compared with RCD cluster A, RCD cluster B had more patients with un-methylated MGMT promoter, chr 19/20 co-gain, wild type IDH, higher-level pathological tissue types and older ages (Fig 3D). The levels of most RCDs were higher in RCD cluster A than in RCD cluster B (Fig 3D), which was consistent with previous results showing that higher RCD levels were detrimental to patients’ prognosis.

The expression of some classical immune checkpoint genes, including CD274, PDCD1, CD247, PDCD1LG2, CTLA4, TNFRSF9, TNFRSF4 and TLR9, was investigated in the RCD clusters, and the results indicated that these genes, except for TLR9, had higher expression in RCD cluster A. Both innate and adaptive immune cells infiltrated at significantly higher levels in RCD cluster A than in RCD cluster B (Fig 3D). The TMB increased in RCD cluster A (Supplementary Fig 1A). The pathway enrichment analysis based on the DEGs between the two RCD clusters revealed that the dysfunction and abnormality of the RCD were significantly related to neutrophil chemotaxis, monocyte chemotaxis, cytokine-cytokine receptor interaction, and collagen degradation (Fig 3E).

We also performed the same analysis method in the TCGA GBM cohort and obtained similar results to those in the TCGA glioma cohort (Supplementary 1B-C, supplementary table 10). Generally speaking, in the context of tumor development, dysregulation of regulated cell death pathways could impact both tumor cells and immune cells, leading to alterations in the immune microenvironment[24]. Some forms of regulated cell death, such as necroptosis and pyroptosis, could induce inflammation and release danger signals (damage-associated molecular patterns - DAMPs)[25, 26]. DAMPs can activate innate immune cells and promote an inflammatory response[27]. This inflammation can influence the recruitment and activation of various immune cells within the tumor microenvironment, ultimately affecting the prognosis of glioma patients.

### Integrated single-cell level analysis of the RCD in glioblastoma

Based on the previous results indicating dysfunction and an abnormal increase in RCD in glioblastoma with bulk-level data, we subsequently investigated RCD using single-cell data to enable a finer scale assessment of the RCD landscape in gliomas. After clustering, dimensionality reduction, and annotation, we present the 12 types of cells and their corresponding numbers (Fig 4A). For each single cell, we estimated the signal pathway activity of RCD using the AUCell method (Supplementary Fig 2). The pathway activity of a cell type was represented by the average of all cells of the same type. Disulfidptosis and immunogenic cell death had higher AUCell scores compared to other RCD types (Fig 4B-D). Notably, disulfidptosis exhibited higher signal levels in endothelial and pericyte cells than in other cell types. The dominant RCD type of each single cell was defined as the type of RCD with the highest level, and we observed a significant proportion of cells with disulfidptosis and immunogenic cell death as the dominant RCD (Fig 4E).

**Fig 4.**
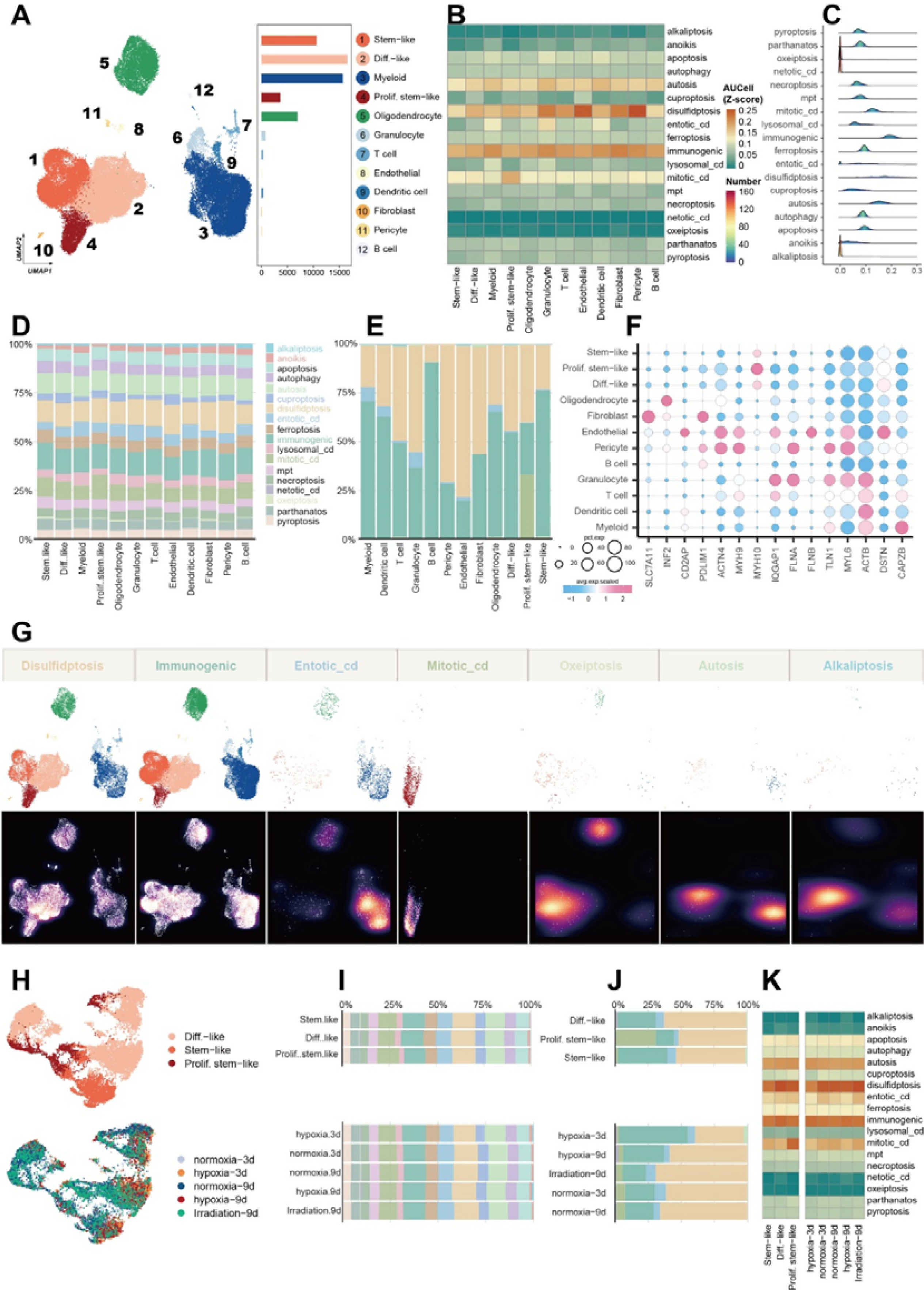
Integrated single-cell level analysis of the RCD in glioblastoma. A. UMAP plot showing that the 12 types of cells and their corresponding numbers were obtained after clustering, dimensionality reduction, and annotation. B. The AUCell of the 18 RCD signatures in the 12 cell types. The pathway activity of a cell type was represented by the average of all cells of the same type. C. The ridge plot showing the distribution of the RCD profile in the single-cell level. D. Percentage of the AUCell scores of the different RCDs in 12 cell types. E. For a single cell, we defined the dominant RCD type of this cell as the type of RCD that has the highest level of its RCD. This plot showing the percentage of the dominant RCD types in 12 cell types. F. Core gene expression of disulfidptosis across defined cell clusters. Bubble size is proportional to the percentage of cells expressing a gene and color intensity is proportional to average scaled gene expression. G. UMAP view of dominant RCD (top) and cell density (bottom) displaying the RCD profile distribution across the different cell types. High relative cell density is shown as bright magma. H. UMAP view of cell types with different stress interventions. I. The percentage of the RCD in different cell types and in different stress interventions. J. The percentage of the dominant RCD in different cell types and in different stress interventions. K. The profile of the 18 RCD in different conditions and cell types.

Given the previous results showing an absolute predominance of disulfidptosis in glioblastoma, we investigated the core disulfidptosis genes in different cell types. SLC7A11 was highly expressed in fibroblast cells, while MYH10 showed high expression levels in malignant cells, especially in Prolif.stem-like cells (Fig 4F). We found that mitotic cell death was the dominant RCD type in almost all the Prolif.stem-like cells. The UMAP view of cells with the dominant RCD (top) and cell density (bottom) displayed the RCD profile distribution across the different cell types (Fig 4G). Furthermore, we investigated the substantial changes in the different RCD cell landscape when exposed to external intervening factors including hypoxia and irradiation (Fig 4H-K, supplementary Fig 3,4).

As the duration of hypoxia increased, the proportion of cells in which disulfidptosis predominated gradually increased, reaching levels similar to controls, while immunogenic cell death gradually decreased to baseline levels (Fig 4J). Overall, we explored the unique landscape of RCD in glioblastomas from a single-cell perspective and found that disulfidptosis and immunogenic cell death appeared to play more important roles in glioblastomas than other RCD types. Additionally, it is worth noting that in the presence of external environmental stimuli, the transformation of these two modes of cell death may be a potentially possible mechanism used by cells to adapt to external stresses.

### Development and validation of a novel RCD-related gene pair signature

The prognosis of glioblastoma is poor, with an overall survival rate remaining relatively low. Even with aggressive treatment, the median survival is typically around 12-16 months[28]. The results presented above suggested that RCD had a significant impact on patients with glioblastoma. Therefore, the purpose of this study is to establish a reliable RCD-related model that can be used in the clinic to accurately predict a patient’s prognosis, quality of life, and response to treatment. This model aims to assist doctors in achieving accurate treatment for their patients. The framework for constructing the scoring system is displayed in Fig 5A.

**Fig 5.**
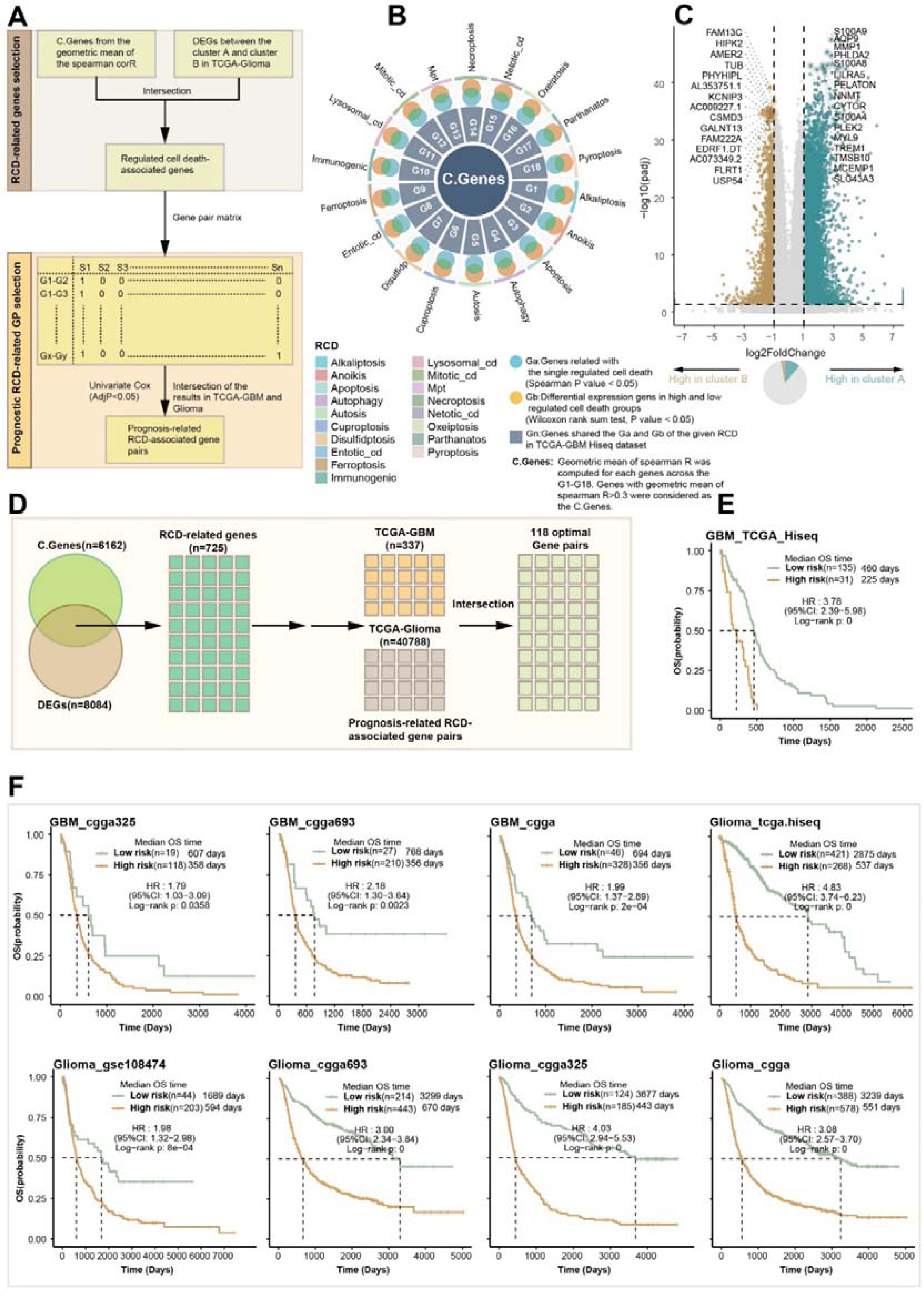
Development and validation of a novel RCD-related gene pair signature. A. The workflow of the construction the RCD.GP scoring system. B. Identification of the C.Genes in TCGA GBM cohort. C. Volcano plot showing the DEGs between the RCD cluster A and RCD cluster B. The process of the obtaining the 118 optimal gene pairs from the DEGs and the C.Genes. E. K-M curves significant difference of the prognosis between the high RCD.GP score subgroup and low RCD.GP score subgroup. The log-rank sum test p < 0.0001. F. The K-M curves in different glioma cohorts shown the same trend, indicating that the patients with high RCD.GP score had worse OS than those with low RCD.GP score.

With calculated geometric spearman’s correlation, we identified 6162 C.Genes related RCD. Detailed descriptions of this process are provided in the methods section (Fig 5B). Additionally, 8084 genes with a p-value less than 0.05 and |log2FC| greater than 1 were identified as DEGs between distinct RCD patterns (Fig 5C). The intersection of C.Genes and DEGs resulted in the final set of RCD-related genes (n = 725). After removing missing genes in most validation datasets, we constructed 639 RCD-related gene pairs. Among the 203,841 gene pairs based on these 639 genes, 377 gene pairs in TCGA-GBM and 40,788 gene pairs were identified as prognostic RCD-related gene pairs using univariate Cox regression (Fig 5D, supplementary Fig 5A, p < 0.05). Finally, 118 gene pairs, including 102 genes, were selected as the optimal gene pairs for constructing the scoring system, named RCD.GP score (Supplementary table 11). In the TCGA-GBM HiSeq dataset, GBM patients with high RCD.GP risk scores exhibited significantly worse prognosis than those with low RCD.GP risk scores (Fig 5E, HR:3.78, 95%CL:2.39-5.98, Log-rank p < 0.0001). This conclusion was also validated in other independent datasets, including GBM_CGGA325, GBM_CGGA693, GBM_CGGA, GBM-GSE108474, Glioma-TCGA, Glioma-GSE108471, Glioma-CGGA, Glioma-CGGA325, CGGA693-Glioma, LGG-CGGA, LGG-CGGA325, LGG-CGGA693, LGG-GSE10611, LGG-GSE108474 and LGG-TCGA (Fig 5F, supplementary Fig 5A, supplementary table 12). The RCD.GP score demonstrated favorable performance in terms of C-index, 1-year AUC, 3-year AUC and 5-year AUC across all 16 datasets (Supplementary Fig 5B). Multivariate Cox regression analysis in TCGA-GBM, GBM-CGGA325, GBM-CGGA693 and GBM-CGGA cohort revealed that the RCD.GP risk score was an independent prognostic factor for glioblastoma patients (Supplementary Fig 5C). The meta-analysis further confirmed the comprehensive and integrated HRs for RCD.GP scores (Supplementary Fig 5D-E). Overall, the RCD.GP score represents a robust and powerful model capable of accurate prognostic predictions for glioma patients. It can aid in early detection, personalized treatment, treatment planning, decision-making, protective management and patient empowerment.

### Glioma with high RCD.GP score possesses strongly malevolent biology and activated immune characteristics

The tumour microenvironment plays a crucial role in tumor growth, immune surveillance, immune escape, and response to therapy[29]. Immune checkpoints are molecules on immune cells that regulate the immune response, preventing excessive activation and tissue damage[30]. However, tumors can hijack these checkpoints to suppress immune responses and avoid immune destruction. Inhibitory immune checkpoint molecules, such as PD-1, PD-L1, and CTLA-4, are frequently expressed within the tumor microenvironment and can dampen immune responses against cancer cells. Our investigation revealed that glioblastomas with high RCD.GP scores highly expressed immune checkpoint genes, including CD274, CD247 and so on (Fig 6A). As the immune microenvironment consists of various immune cells, including lymphocytes, macrophages, dendritic cells, and myeloid-derived suppressor cells (MDSCs) which can infiltrate the tumor site and interact with cancer cells, influencing tumor progression, we assessed the infiltration of the immune cells, and found that higher RCD.GP risk scores correlated with increased infiltration of cells such as M0 macrophages, neutrophils, and Tregs (Fig 6A). Similar results were observed in the CGGA dataset (Supplementary Fig 6A). To further confirm the association of RCD.GP scores with malignant biology and tumor-associated immunity modulation in glioblastoma, we performed GSVA with the cancer hallmark signature database, KEGG database, REACTOME database and GO BP database in TCGA-Glioma. The high RCD.GP scores were associated with activated malignant pathways and immune-related signals, such as epithelial mesenchymal transition, complement and coagulation cascades, cytokine-cytokine receptor interaction, and complement cascade (Fig 6B-C, supplementary Fig B-D). The enrichment analysis with GO, KEGG and REACTOME pathways further validated the activation of immune-related signals and malignant pathways in the high RCD.GP score subgroup (Fig 6D, supplementary Fig 6E-F). The anti-tumor immune cycle, representing the steps and interactions involved in mounting an effective immune response against tumors[30], exhibited higher signals in the high risk score subgroup, indicating an activated anti-tumor immune response.

**Fig 6.**
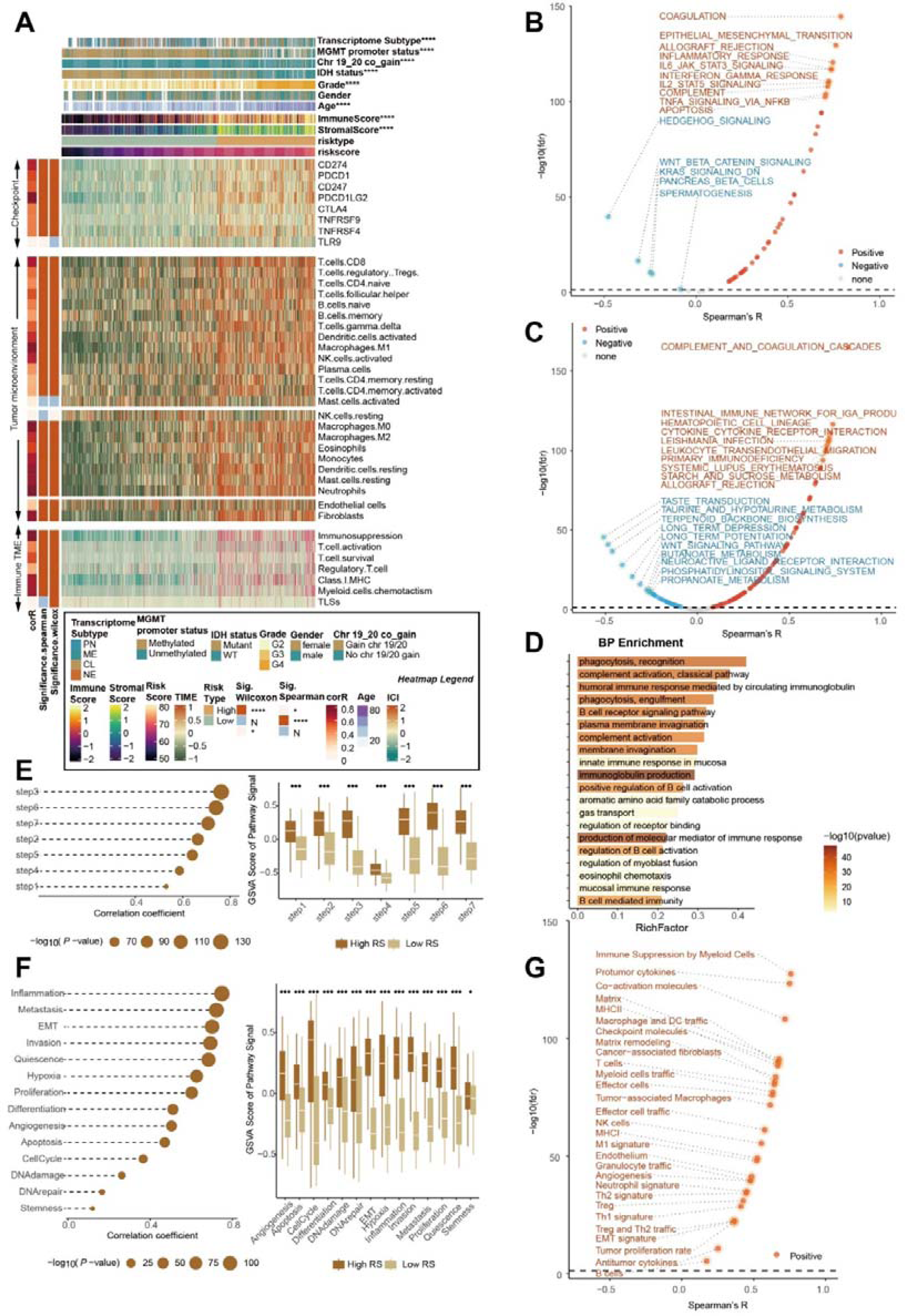
Glioma with high RCD.GP score possesses strongly malevolent biology and activated immune characteristics. A. The heat map of the relationship of the RCD.GP score and clinical indexes, immune checkpoint genes, infiltration of the immune cells and immune microenvironment function in TCGA glioma cohort. B-C. The spearman’s correlation between the RCD.GP score and the GSVA score of the tumor hallmark signatures (B) and functional pathway signatures in KEGG database (C). E-F. The plot showing the relationship between the RCD.GP score and the anti-tumor cycle (E), and single cell state from the cancerSEA (F). The spearman’s correlation and the Wilcoxon rank sum test were performed. “*”, “**”, “***”, and “****” represented that the p value < 0.05, 0.01, 0.001, and 0.0001. G. The spearman’s correlation between the RCD.GP score and the 29 signatures related with the tumor microenvironment.

We also estimated the correlation between a number of immune-related scores [31] (n = 92) and RCD.GP scores, and the results indicated a positive relationship between most immune-related scores and the RCD.GP score (Supplementary Fig 6G). Although patients with higher risk scores had worse prognoses, we investigated the features of the tumor microenvironment using signatures from the cancer single-cell states atlas [32] (n = 14) and the previous literatures (n = 29)[33]. The GSVA results revealed that factors promoting intense malignant progression of tumors, such as EMT, inflammation, metastasis, immune suppression by myeloid cells, pro-tumor cytokines, and tumor proliferation, were significantly higher in the high-risk group than in the low-risk group (Fig 6F-G, supplementary Fig 6H). One possible explanation for the apparent contradiction is that the tumor microenvironment, characterized by factors such as inflammation, immune cell infiltration, hypoxia, and nutrient deprivation, can influence the levels of cell death in tumors. Certain aspects of the tumor microenvironment could promote cell death, while others could protect tumor cells from undergoing programmed cell death. In glioblastoma, both might be at a high level, leading to an imbalance that favors increased cell death but does not necessarily result in tumour shrinkage. This imbalance could be due to the rapid and uncontrolled proliferation of tumor cells, leading to an increased number of cells requiring elimination.

### Assessment of the capability of the RCD.GP score for the immunotherapy response

Considering the impressive association of the RCD.GP score with immune-related characteristics, we hypothesized that the RCD.GP score could be strongly linked to the response to immunotherapy. To investigate this, we collected some signatures related to immunotherapy response and found that the RCD.GP score positively correlated with the GSVA scores of these signatures (Fig 7A).

**Fig 7.**
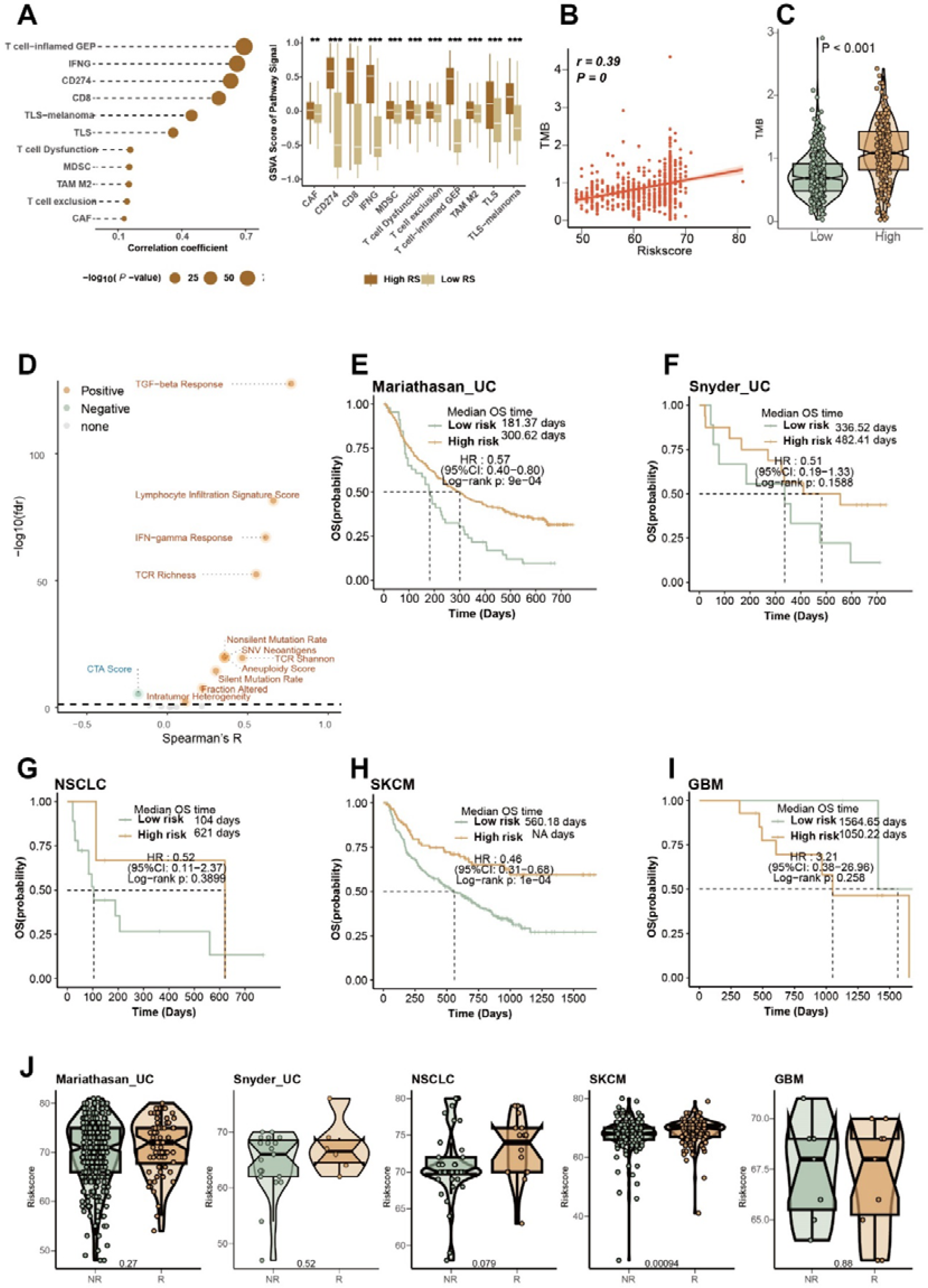
Assessment of the capability of the RCD.GP score for the immunotherapy response. A. The relationship of the RCD.GP score and the GSVA score of the signatures related with the immunotherapy response. The spearman’s correlation analysis and the Wilcoxon rank sum test were performed. “*”, “**”, “***”, and “****” represented that the p value < 0.05, 0.01, 0.001, and 0.0001. B. The spearman’s correlation between the RCD.GP score and the tumor mutation burden. C. The difference of the TMB between the high RCD.GP score subgroup and low RCD.GP score subgroup. The Wilcoxon rank sum test was used. D. The spearman’s correlation of the RCD.GP score and the immune scores related with the immunotherapy response. E-I. The K-M curves showing the difference of the prognosis between the high RCD.GP score subgroup and the low RCD.GP score subgroup in UC, SKCM, NSCLC and GBM. J. The difference of the RCD.GP score between the NR patients and R patients. NR: not response to immunotherapy response. R: response to immunotherapy response.

Previous studies have demonstrated that tumors with high TMB are more likely to respond to immunotherapy due to enhanced recognition and targeting by the immune system [34, 35]. We observed a positive correlation between the RCD.GP score and the TMB (Fig 7B, R=0.39, p < 0.001). Furthermore, the TMB was higher in the high-risk score group compared to the low-risk score group (Fig 7C, Wilcoxon rank sum test p < 0.001). Other immunotherapy response-related scores, such as TCR richness, TGF-beta response, and lymphocyte infiltration signature score, also showed positive associations with the RCD.GP score (Fig 7D, supplementary table 13). In the high-risk score group, most of these scores were higher than those in the low-risk score group (Supplementary Fig 7).

To further verify the predictive efficacy of the RCD.GP score in immunotherapy response, we analyzed multiple cohorts with immunotherapy treatment. Strikingly, the higher RCD.GP score group exhibited better prognosis and immunotherapy response in patients with UC, NSCLC and SKCM, while intriguingly, the K-M curves exhibited the opposite trend in patients with glioblastoma (Fig 7E-J, supplementary table 14). Overall, our study suggests that patients with a high RCD.GP score may have a higher potential to benefit from immunotherapy treatment.

### Identification of SLC43A3 as novel and potential RCD-related oncology target

In this study, we present a novel framework for screening important features (Fig 8A). Differentially expressed gene analysis was performed among GBM, LGG and normal brain cortex (Supplementary Fig 8A-C), leading to the identification of 3260 genes as DEGs across all groups (Fig 8B). Among these, 725 genes were identified as RCD-related genes in glioma, and 1660 genes were associated with the prognosis of the glioma patients (Fig 8A-B). From this analysis, 60 genes were identified as RCD-related prognostic DEGs (Fig 8B, supplementary Fig 8D). Using a machine learning framework on TCGA-Glioma cohort and TCGA-GBM cohort, we generated some candidate genes (Fig 8C-D).

**Fig 8.**
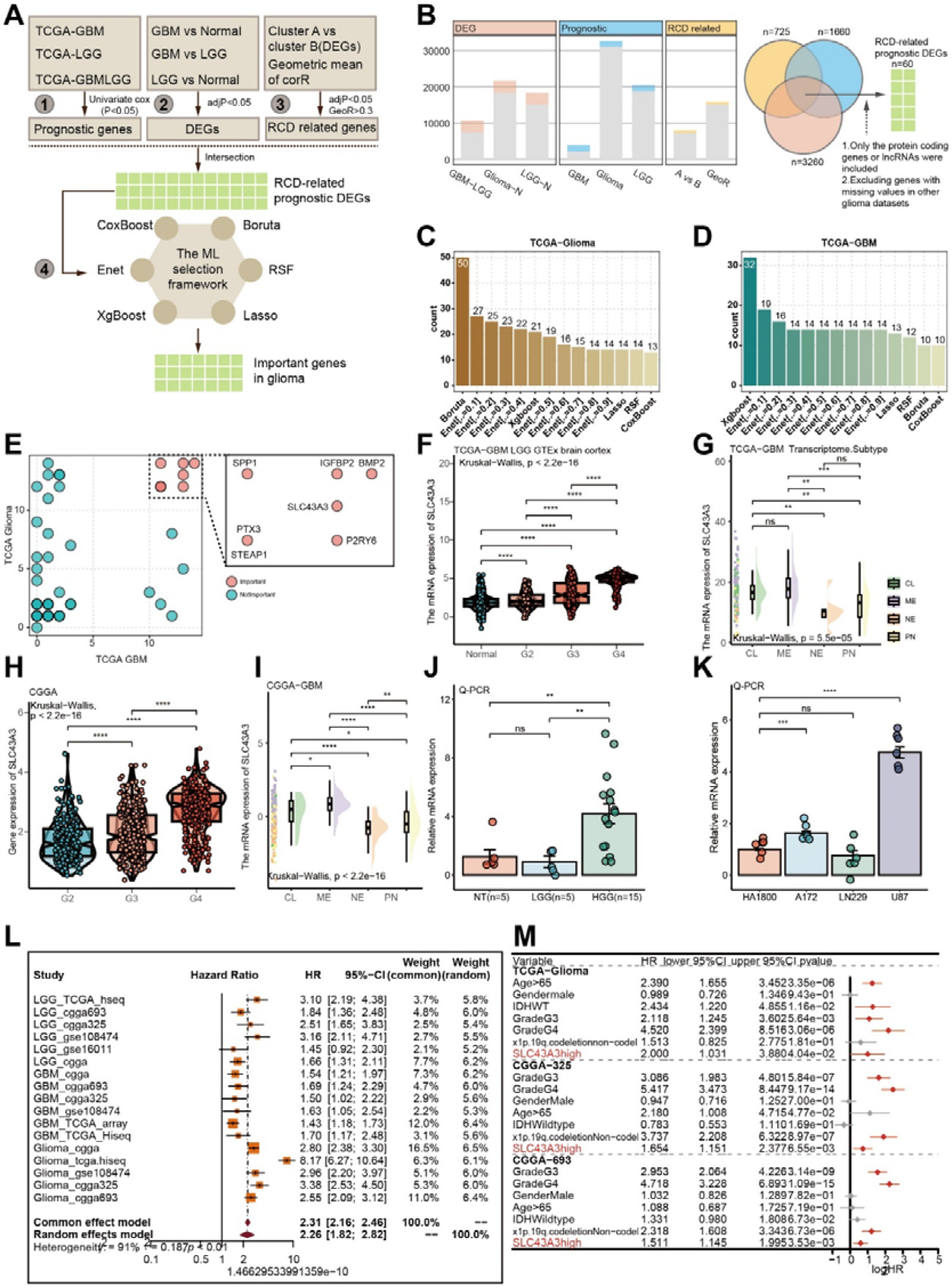
Identification of SLC43A3 as novel and potential RCD-related oncology target. A. The framework of screening the important features with machine learning methods. B. The numbers of the DEGs, RCD related genes, and prognostic candidates. C-D. The number of the important genes screened by different machine learning methods with different parameters in TCGA-Glioma cohort (C) and TCGA-GBM cohort (D). E. The number of times each genes was screened out by the machine learning framework in TCGA-Glioma cohort and TCGA-GBM cohort. F. The profile of the SLC43A3 in different grade glioma and normal brain cortex tissues in TCGA and GTEx datasets. Wilcoxon rank sum test was used for assessing the difference between the two subgroups. “*”, “**”, “***”, and “****” represented that the p value < 0.05, 0.01, 0.001, and 0.0001.G. The difference of the SLC43A3 expression among the different transcriptomic subtypes in TCGA-GBM cohort. H-I. The expression of the SLC43A3 in different glioma grades and different transcriptomic subtypes in CGGA cohort. J. The Q-PCR result showing the SLC43A3 highly expressed in HGG compared with LGG and normal brain cortex tissues. NT: normal brain cortex tissue. LGG: low-grade glioma. HGG: high-grade glioma. The Wilcoxon rank sum test was utilized. K. Q-PCR of the SLC43A3 relative expression in different glioma cell lines. The Wilcoxon rank sum test was utilized for confirming the difference. L. The meta analysis provided a comprehensive HR of the RCD.GP score. M. Multivariate Cox regression analysis of the SLC43A3 was performed in TCGA-Glioma, CGGA-325 and CGGA-693 cohort.

The genes that were repeatedly screened out more than 10 times in both TCGA-GBM and TCGA-Glioma cohorts were identified as the most important genes for glioma patients (Fig 8E, supplementary table 15). These genes included BMP2, IGFBP2, SPP1, SLC43A3, P2RY6, PTX3, and STEAP1. Among these, the role of SLC43A3 in tumors, especially gliomas, had hardly been reported, while the functions of the other genes in tumors were widely documented [36-41]. Interestingly, the expression of SLC43A3 increased with the pathological grade and was most highly expressed in the ME transcriptional subgroup, which has been associated with a malignant process and shorter median survival time in glioblastoma patients (Fig 8F-I). We also comprehensively investigated the distribution of SLC43A3 in different clinical subgroups, revealing a potential relationship between SLC43A3 expression and poor clinical prognosis (Supplementary Fig 9A-M).

To validate the findings, we assessed SLC43A3 expression in high-grade gliomas compared to normal brain cortex using q-PCR (Fig 8J). Transcriptomic data from the CCLE project showed higher mRNA levels of SLC43A3 in most glioblastoma cells compared to LGG cells (H4) (Supplementary Fig 9N). The q-PCR experiment with glioblastoma cell lines further confirmed higher expression of SLC43A3 in glioblastoma cell lines compared to astrocyte cell lines (Fig 8K). Prognostic analysis using multiple glioma datasets revealed that high expression of SLC43A3 was associated with worse overall survival (Supplementary Fig 10A-C). Meta-analysis and multiple variate Cox regression analysis provided strong evidence that SLC43A3 was a risky and independent prognostic factor for predicting glioma patients’ outcomes (Fig 8L-M).

In conclusion, we have provided a novel framework for screening important features, identified seven potential important therapeutic targets, and presented solid evidence supporting an oncogenic role of RCD-related SLC43A3 in glioblastoma.

## Discussion

Glioblastoma is the most common and deadliest type of brain tumor in adults, accounting for approximately 15% of all primary brain tumors. It originates from glial cells in the brain, and is characterized by uncontrolled cell growth, invasion into surrounding tissues, and resistance to cell death. Despite advancements in treatment, the prognosis for glioblastoma remains poor, with a median survival time of about 12 to 16 months. Its highly invasive nature and resistance to therapies make it challenging to treat, emphasizing the urgent need for more accurate predictive tools to improve overall survival rates, symptom management, and patients’ quality of life.

Previous predictive signatures for glioblastoma patient prognosis mainly relied on individual gene expression, disregarding the interactions or synergistic relationships between genes. In contrast, predictive models based on gene pairs offer several advantages including increased predictive power, enhanced interpretability and transferability to different datasets or experimental conditions[42]. Here, we comprehensively investigated the RCD landscape of glioblastoma from both bulk and single-cell perspectives. We developed a robust and accurate RCD-related gene pair signature that holds potential for transferability across different datasets. Additionally, we provided a framework for screening out relatively core genes and identified SCL43A3 as a potential therapeutic oncology target.

Our results revealed a complex and multifaceted relationship between RCD and glioblastoma. The elevated levels of almost all RCD in glioblastoma were associated with worse prognosis, strongly correlated with malignant biology processes and immune microenvironment dysfunction. Cancer cells often exploit immune checkpoint pathways to evade immune surveillance and promote tumor growth [43]. They upregulate immune checkpoint molecules on their surface or within the tumor microenvironment, leading to the suppression of anti-tumor immune responses. Glioma with higher levels of RCD exhibited increased expression of immune checkpoint genes, indicating potential suppression of anti-tumour immunity. Combining therapies that induce regulated cell death with immune checkpoint inhibitors have shown promising results in preclinical and clinical studies[44-48]. Inducing immunogenic forms of cell death can enhance the immunogenicity of tumors and improve the efficacy of immune checkpoint blockade by promoting the activation of anti-tumor immune responses[49]. Signaling pathways promoting malignant biological processes such as JAK-STAT signaling pathway, TNF signaling pathway and NF-kappa B signaling pathway, were demonstrated to be extraordinarily related with the high level of RCD. However, we also observed enhanced anti-tumor immune responses, such as the increased infiltration of anti-tumor immune cells and the activated anti-tumor immune signals like IL-17 signaling pathway, cytokine-cytokine receptor interaction and neutrophil chemotaxis which might be a result of the dysregulated tumor microenvironment promoting both cell death and tumor protection mechanisms. Further investigation is required to understand the complexity of molecular mechanisms underlying this association and its therapeutic implications.

Single-cell analysis suggested a significant role for disulfidptosis in glioblastoma. Disulfidoptosis, also known as disulfide bond deficiency, is a condition characterized by an impaired formation or maintenance of disulfide bonds [23]. Disulfide bonds are important for the stability and proper folding of proteins within cells. Disulfide bond deficiency can potentially impact various cellular processes that are relevant to tumorigenesis. One possible link is the role of disulfide bonds in protein folding and cell signaling. Proteins involved in cell growth regulation, regulated cell death, and DNA repair often require proper disulfide bond formation to function correctly. Disruptions in these processes can contribute to the development and progression of tumors. The expression of SLC7A11, a core gene of disulfidoptosis, was highly expressed in fibroblast cells, while MYH10, another core gene, showed high expression in malignant cells, particularly Prolif.stem-like cells. This suggests MYH10 as a potential therapeutic target for glioma. Nonetheless, the relationship between disulfidoptosis and glioblastoma is complex, requiring further investigation.

The RCD.GP score proved to be a robust and promising biomarker for predicting clinical outcomes and immunotherapy response in glioma patients. Traditional prediction models based on absolute gene expression values may be influenced by noise and biological variability, leading to less reliable predictions. In contrast, our model, incorporating gene pairs, demonstrates improved dtability and robustness by accounting for variability in gene expression data. In 16 cohorts of glioma patients, the RCD.GP score consistently exhibited superior reliability and generalizability after validation and optimization. The validation metrics including, K-M curves, C-index, 1-year AUC, 3-year AUC, 5-year AUC, meta-analysis results, and multivariate Cox regression analysis, unequivocally support the enhanced predictive power of the RCD.GP score system.

Patients with glioma and high RCD.GP scores experienced shorter overall survival times, which were strongly associated with malignant biological processes, including coagulation, epithelial-mesenchymal transition, and inflammatory response. Additionally, the subgroups with high RCD.GP risk scores displayed an increased profile of immune checkpoint genes, enhanced infiltration of the immune cells, and elevated levels of pro-tumor signaling pathways. Immunotherapy has demonstrated promising results in various cancers, such as melanoma, lung cancer, bladder cancer, and some types of lymphomas and leukemia[50-54]. Our study also revealed a significantly positive correlation between the RCD.GP score and immunotherapy response indexes, including CD8, MDSC, TLS, TMB and TGF-beta response.

Interestingly, while patients with high RCD.GP scores showed improved OS and enhanced immunotherapy response in various cancer types like UC, NSCLC and SKCM, a different trend was observed in GBM patients treated with anti-PD-1 therapy. The group with higher RCD.GP scores exhibited a worse prognosis and impaired immunotherapy response. Despite the promising results of immunotherapy in treating other cancers, immune checkpoint inhibitors have shown limited efficacy in glioblastoma compared to traditional treatment modalities such as radiotherapy, chemotherapy and surgery[55]. This may be attributed to the intricate interplay between multiple immune-related mechanisms affecting glioblastoma progression and the influence of RCD in promoting malignant progression. As such, exploring combination approaches, such as using different immunotherapies in conjunction with RCD inhibition, chemotherapy, radiation therapy, or targeted therapies, holds potential for further improving treatment outcomes in glioblastoma patients.

Seven genes including BMP2, IGFBP2, SPP1, SLC43A3, P2RY6, PTX3 and STEAP1 were identified as potential therapeutic targets through a machine learning framework designed to identify the most important features. Each of these genes plays a unique role in tumorigenesis and tumor progression. BMP2 is a multifaceted gene that can exhibit both tumor-promoting and tumor-suppressive effects in different types of cancers. It can stimulate cell proliferation, angiogenesis, and metastasis in certain contexts, while also inducing cell cycle arrest, apoptosis (programmed cell death), and differentiation of cancer cells, leading to tumor growth inhibition[36]. IGFBP2 has been implicated in promoting tumor growth and progression in several types of cancer [37]. SPP1, a glycoprotein with diverse functions, is involved in various biological processes such as cell adhesion, migration, immune regulation, and tissue remodeling. Its ability to stimulate cell proliferation, survival, and angiogenesis, facilitating tumor formation [38, 56]. P2RY6 and STEAP1 are considered tumor suppressor genes in certain cancer types [41, 57]. PTX3 is known to enhance tumor cell proliferation, survival, invasiveness, and angiogenesis by modulating various signaling pathways in different tumor types, including breast cancer, lung cancer, ovarian cancer, and glioblastoma [58]. SLC43A3, a protein-coding gene that encodes a transporter involved in amino acid transport, has been associated with fatty acid flux, nucleotide metabolism and DNA repair [59-61]. Its role in cancer development and progression is not yet fully understood. Our focused investigation on SLC43A3 revealed a RCD-related oncogenic potential in glioma through systematic analysis of its expression profile and prognostic value in multiple datasets and q-PCR experiments. Overall, the identification of these seven genes as potential therapeutic targets opens up new avenues for future research and the development of targeted therapies for glioma treatment. Further studies are warranted to elucidate the precise molecular mechanisms and potential We acknowledge that this study has some limitations. Firstly, the cohorts with immunotherapy included only four cancer types, and to draw a more accurate and broadly generalized conclusion, data from a wider variety of cancer cohorts with immunotherapy should be included in the analysis. Secondly, the single-cell analysis revealed the core role of disulfidoptosis in glioblastoma, but only a rudimentary analysis was conducted. To comprehensively elaborate its role, further investigations involving more systematic, comprehensive, and advanced analysis and experiments are necessary. Thirdly, although we identified seven candidate genes for therapy, we only validated the expression profile of SLC43A3 in tissues and cancer cell lines using q-PCR. To gain a deeper understanding of the potential and insightful mechanisms of SLC43A3, additional experiments need to be conducted. Finally, this study utilized retrospective data without prospective clinical trials data for validating the superior reliability and generalizability of the RCD.GP score. To further validate the performance of the RCD.GP score, prospective clinical trials data should be incorporated in future studies.

## Conclusions

In conclusion, we have conducted a comprehensive investigation of the RCD landscape in glioblastoma, exploring both bulk and single-cell aspects. Our study resulted in the development of a robust and accurate RCD-related gene pair signature, which holds promise for potential clinical applications. Moreover, we established a framework for identifying relatively core genes and successfully identified SCL43A3 as a potential therapeutic target in oncology. Furthermore, our findings highlight the significance of the RCD.GP score as a predictor for adverse clinical outcomes and impaired immunotherapy response in glioblastoma patients. These insights have the potential to improve patient management and treatment decisions.

## Authors’ contributions

Wei Zhang designed the project, analyzed the data and wrote the manuscript; Xuejun Li and Nian Jiang conceived the project, analyzed the data and revised the manuscript. Hongyi Liu, and Luohuan Dai participated in experiment design and provided a lot of valuable advice. Yihao Zhang, Hongwei Liu and Ruiyue Dang helped collect and visualise the data. Abraham Ayodeji Adegboro made significant contributions to the linguistic refinement and enhancement of this article. All authors contributed to the article and approved the submitted version.

## Funding

This work was supported by the National Natural Science Foundation of China (No.81770781 and No.82270825), Special funds for innovation in Hunan Province (No2020SK2062), Hunan Provincial Natural Science Foundation of China (No. 2022JJ40793), and High talent project of Hunan Province (No. 2022WZ1031).

## Availability of data and materials

No new data except from the q-PCR results was generated as part of this study. All data used in this study were sourced from the public domain online. Additionally, all key codes utilized in this study were available on Github (https://github.com/zwxiangya/RCD.GPscore).

## Acknowledgements

The authors would like to express their gratitude to the following datasets: TCGA, CGGA, GTEx, GSE16011 and so on, for their availability and contribution to this study. The authors also thanked the website (BioRender.com) for helping create the graphic abstract.

## Ethics approval and consent to participate

The study was approved by the ethics committee of Xiangya Hospital, and the written informed consent was obtained from all patients.

## Notes

### Competing Interest Statement

The authors have declared no competing interest.

